# Extracellular RNA signatures of mutant KRAS(G12C) lung adenocarcinoma cells

**DOI:** 10.1101/2022.02.23.481574

**Authors:** Reem Khojah, Roman E. Reggiardo, Mehmet Ozen, Sreelakshmi Velandi Maroli, David Carrillo, Utkan Demirci, Daniel H. Kim

**Affiliations:** Department of Biomolecular Engineering, University of California Santa Cruz, Santa Cruz, CA 95064, USA; Bio-Acoustic MEMS in Medicine (BAMM) Laboratory, Department of Radiology, Stanford University School of Medicine, Palo Alto, CA 94305, USA; Canary Center at Stanford for Cancer Early Detection, Stanford University School of Medicine, Palo Alto, CA 94305, USA; Department of Molecular, Cell and Developmental Biology, University of California Santa Cruz, Santa Cruz, CA 95064, USA; Institute for the Biology of Stem Cells, University of California Santa Cruz, Santa Cruz, CA 95064, USA; Genomics Institute, University of California Santa Cruz, Santa Cruz, CA 95064, USA; Center for Molecular Biology of RNA, University of California Santa Cruz, Santa Cruz, CA 95064, USA

## Abstract

Extracellular RNAs (exRNAs) are actively secreted from cells in membrane-bound extracellular vesicles (EVs). Diverse classes of RNAs are secreted as exRNAs, including messenger RNAs (mRNAs), long noncoding RNAs (lncRNAs), and transposable element RNAs (TE RNAs). However, the full composition and clinical utility of exRNAs secreted in response to oncogenic signaling are unknown. Here we use both affinity- and nanofiltration-based EV isolation approaches to show that mutant KRAS(G12C) signaling results in the secretion of specific lncRNAs, TE RNAs, and mRNAs, some of which are prognostic for lung adenocarcinoma (LUAD) patient survival. We found that inhibition of KRAS(G12C) signaling broadly reprograms the noncoding transcriptome, as evidenced by a substantial increase in TE RNA secretion. KRAS(G12C) inhibition also increased the abundance of secreted lncRNAs and retained intron-containing transcripts, while decreasing the mRNA content of EVs. Oncogenic KRAS(G12C) signaling was required for the secretion of mRNAs from a set of 20 genes that are significantly associated with unfavorable clinical outcomes in LUAD. Our study suggests that both coding and noncoding RNAs that are secreted in EVs may serve as KRAS(G12C)-specific signatures for diagnosing lung cancer.

## INTRODUCTION

More than 75% of the human genome is transcribed into RNA, most of which is noncoding^1^. These noncoding RNAs include long noncoding RNAs (lncRNAs), transposable element RNAs (TE RNAs), microRNAs, and other classes of RNAs that are involved in a diverse array of biological processes^2-6^. A subset of lncRNAs and TE RNAs are regulated by fundamental signaling pathways that are dysregulated during cancer formation, such as the RAS signaling pathway^7,8^. Moreover, these noncoding RNAs are preferentially secreted in extracellular vesicles (EVs)^8^, which are membrane-bound, nanometer-sized vesicles that protect extracellular RNAs (exRNAs) from degradation^9^. Many studies have described the biomarker potential of exRNAs for cancer diagnosis^10^, but the clinical utility of mutation-specific exRNA signatures remains unclear.

The RAS family of signaling proteins are highly conserved and mediate multiple cellular functions, including cell proliferation^11^. The KRAS gene is mutated in ~20% of global cancer patients^12^, and the fraction of lung adenocarcinoma (LUAD) cancers with a mutant KRAS allele is even higher: estimates from The Cancer Genome Atlas (TCGA) indicate that ~30% of patients have detectable KRAS mutations^13,14^. Moreover, ~85% of KRAS mutations occur at codons 12, 13, or 61^13^, and mutations at Glycine 12 lock this GTPase in its active, GTP-bound state that enables constitutive oncogenic signaling^11^. Long thought to be undruggable, KRAS(G12C) has been successfully targeted by small molecule inhibitors^15^, including AMG 510, MRTX849, ARS1620, JDQ443, GDC-6036, LY3537982, BI 1823911, and others^16–19^.

Here we report our transcriptomic analysis of exRNA signatures of mutant KRAS(G12C) in LUAD cells using affinity- and nanofiltration-based EV isolation methods. We demonstrate the utility of using a KRAS(G12C)-specific inhibitor to identify KRAS(G12C)-specific exRNA signatures, which exhibit strong concordance with transcriptomic signatures of mutant KRAS(G12C) primary tumor samples from the TCGA LUAD dataset. Comparative analysis of exRNAs from control and KRAS(G12C) inhibitor-treated LUAD cells revealed that KRAS(G12C)-mediated oncogenic signaling is required for the secretion of specific lncRNAs, TE RNAs, and protein-coding RNAs, a subset of which have prognostic potential.

## RESULTS

### KRAS(G12C) inhibition decreases cell viability and EV size

To determine the effects of mutant KRAS(G12C) inhibition on EVs and exRNAs secreted by H358 LUAD cells (**Fig. 1A**), AMG 510 concentrations ranging from 0.1 nM to 10 *μ*M were added to two-dimensional (2D) adherent monolayer and three-dimensional (3D) spheroid cultures to assess cell viability after 72 hours of inhibitor treatment (**Fig. 1B**). 2D and 3D cell cultures retained ~70% and ~40% viability compared to controls, respectively, after treatment with 0.1 *μ*M of AMG 510 (**Fig. 1B**). EVs were then isolated using two orthogonal approaches: affinity-based (exoRNeasy) and nanofiltration-based (ExoTIC: Exosome Total Isolation Chip)^20–23^. Each EV isolation method captured distinct EV populations, as determined by nanoparticle tracking analysis (NTA) (**Fig. 1C**). The ExoTIC platform captured a population of EVs centered at ~155 nanometers (nm) in diameter from control LUAD cells, while capturing several smaller populations of EVs centered at ~120 nm, ~85 nm, and ~60 nm in KRAS(G12C) inhibitor-treated cells (**Fig. 1C**). The exoRNeasy platform captured a slightly larger population of EVs centered at ~272 nm in diameter from control cells, with KRAS(G12C) inhibitor-treated cells again exhibiting a downshift in size of ~35 nm to a population of EVs centered at ~236 nm in diameter (**Fig. 1C**), suggesting that attenuation of oncogenic KRAS(G12C) signaling decreases EV size.

**Fig. 1.**
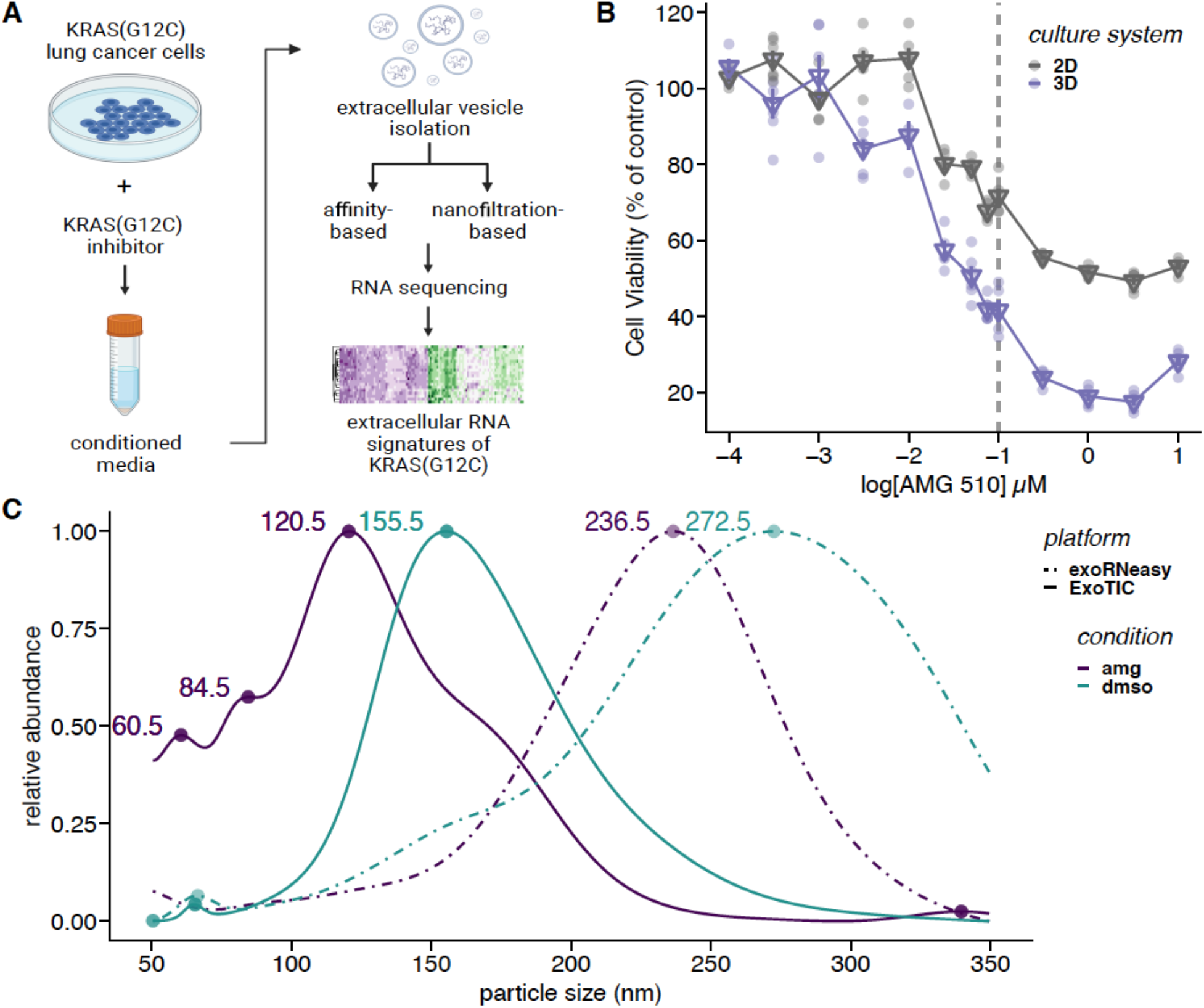
KRAS(G12C) inhibition decreases cell viability and EV size. **A)** Workflow schematic for EV isolation and RNA-seq analysis of KRAS(G12C) exRNA signatures. **B)** Cell viability of 2D and 3D LUAD cells were analyzed after 72 hours of AMG 510 treatment relative to DMSO-treated control cells. Triangular points represent the means calculated for each treatment concentration. **C)** Size distribution of EVs isolated using affinity-based (exoRNeasy) or nanofiltration-based (ExoTIC) isolation platforms, as determined by nanoparticle tracking analysis.

### KRAS(G12C)-dependent exRNA transcriptional landscape

To comprehensively profile the extracellular transcriptome of mutant KRAS(G12C) LUAD cells, we performed RNA sequencing (RNA-seq) using biological triplicates from 4 different conditions: 1) exoRNeasy exRNAs from control LUAD cells (DMSO_2D), 2) exoRNeasy exRNAs from KRAS(G12C) inhibitor-treated LUAD cells (AMG_2D), 3) ExoTIC exRNAs from control LUAD cells, and 4) ExoTIC exRNAs from KRAS(G12C) inhibitor-treated LUAD cells. The smaller EVs isolated using the ExoTIC platform exhibited greater transcriptional complexity than the slightly larger EVs isolated using the exoRNeasy platform, based on the number of different transcripts detected (**Fig. 2A**). KRAS(G12C) inhibitor treatment also led to a consistent and reproducible increase in exRNA transcriptional complexity across both EV isolation platforms (**Fig. 2A**), suggesting that a macro-level hallmark of KRAS(G12C) inhibition is an increase in the diversity of exRNAs secreted from LUAD cells.

**Fig. 2.**
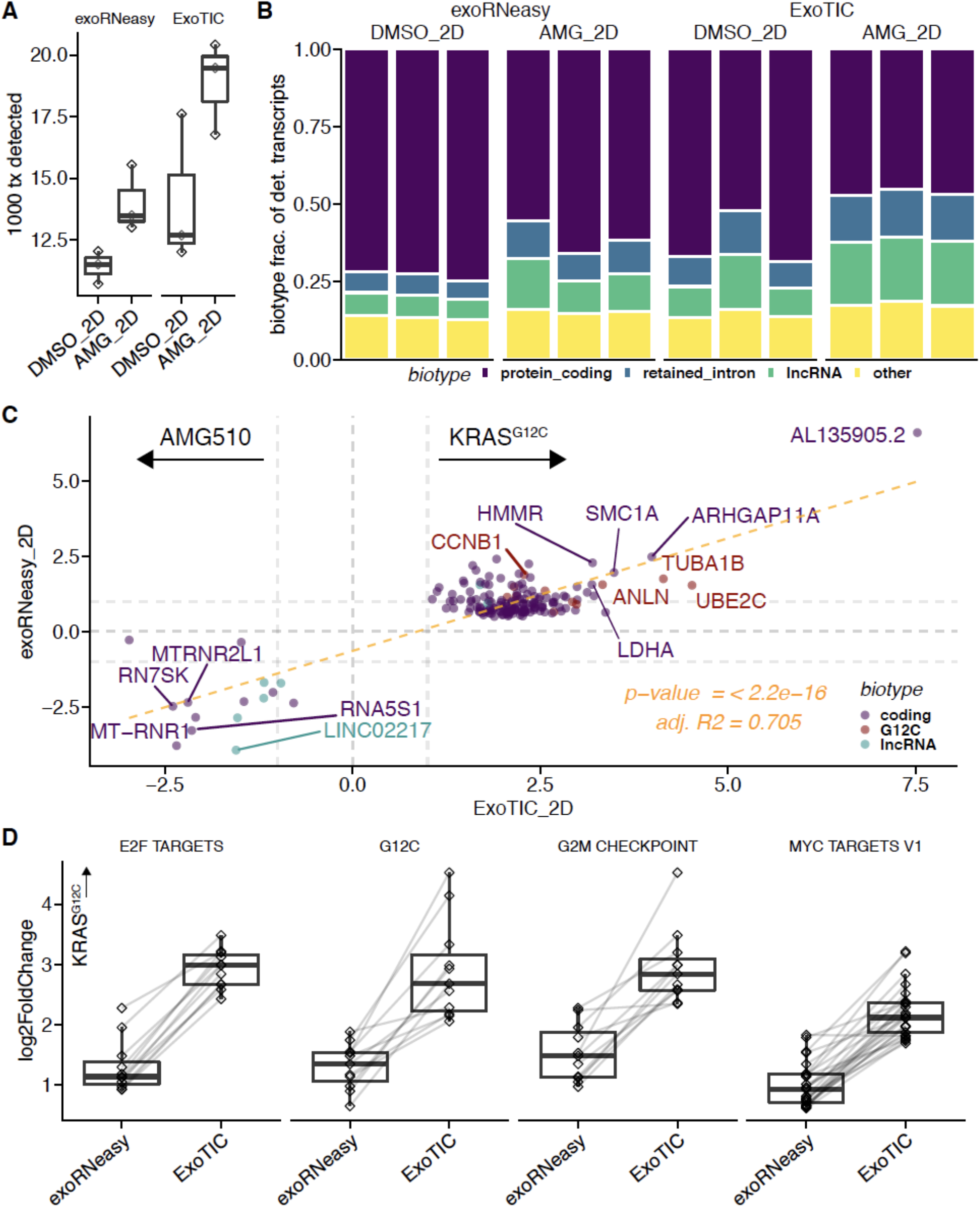
KRAS(G12C)-dependent exRNA transcriptional landscape. **A**) Distribution of the number of transcripts detected above a threshold of 5 normalized counts in both ExoTIC and exoRNeasy platforms. **B**) Stacked bar plot displaying the fraction of detected transcripts annotated as protein-coding, retained intron, lncRNA, or other GENCODE biotypes. **C**) Scatter plot comparing log2-scale fold-changes between AMG and DMSO treatment using the ExoTIC (x-axis) and exoRNeasy (y-axis) platforms. Colors represent GENCODE biotypes lncRNA, protein-coding, or membership in the KRAS(G12C)-induced gene set from Xue *et al* ^24^. **D**) Distribution of log-scale fold-changes of genes included in Hallmark gene sets enriched in both ExoTIC and exoRNeasy platforms, as well as the KRAS(G12C)-induced gene set mentioned above.

LUAD cells with intact KRAS(G12C) signaling primarily secreted protein-coding exRNAs, along with lncRNAs and RNAs containing retained introns (**Fig. 2B**). Upon inhibition of KRAS(G12C) signaling, cells secreted a larger fraction of noncoding RNAs in both exoRNeasy- and ExoTIC-isolated EVs, and for ExoTIC exRNAs, more than half of all secreted exRNAs were noncoding (**Fig. 2B**). Both EV isolation platforms also showed agreement across a subset of secreted, differentially expressed (DE) genes between cells with intact KRAS(G12C) signaling and cells treated with KRAS(G12C) inhibitor (**Fig. S1**), with 64 genes that were found to be significantly enriched in EVs from cells with intact KRAS(G12C) signaling (**Fig. 2C**). Moreover, hallmark genes of mutant KRAS(G12C) signaling^24^ were significantly depleted in EVs isolated from cells treated with KRAS(G12C) inhibitor, providing transcriptomic evidence for the inhibitory role of AMG 510 treatment on KRAS(G12C) signaling (**Fig. 2C**). DE exRNAs from both EV platforms also demonstrated strong agreement using Gene Set Enrichment Analysis (GSEA), with overlapping enrichment of several hallmark gene sets in cells with intact KRAS(G12C) signaling: MYC TARGETS V1, E2F TARGETS, and G2M CHECKPOINT (**Figs. 2D and S1**)^25^. These results show that oncogenic KRAS(G12C) signaling is required for the secretion of a common set of functionally related exRNAs.

### KRAS(G12C) inhibition increases secretion of noncoding RNAs in EVs

We next examined how KRAS(G12C) inhibition affects the composition of secreted exRNAs. 347 genes were significantly enriched in EVs from cells treated with KRAS(G12C) inhibitor (**Fig. 3A**), in contrast to the 1,787 genes significantly enriched in EVs from cells with intact KRAS(G12C) signaling (**Fig. S1**), suggesting that KRAS(G12C) inhibition leads to a decrease in protein-coding mRNA secretion. However, KRAS(G12C) inhibition led to a significant increase in lncRNA secretion when compared to cells with intact KRAS(G12C) signaling (**Fig. 3B**).

**Fig. 3.**
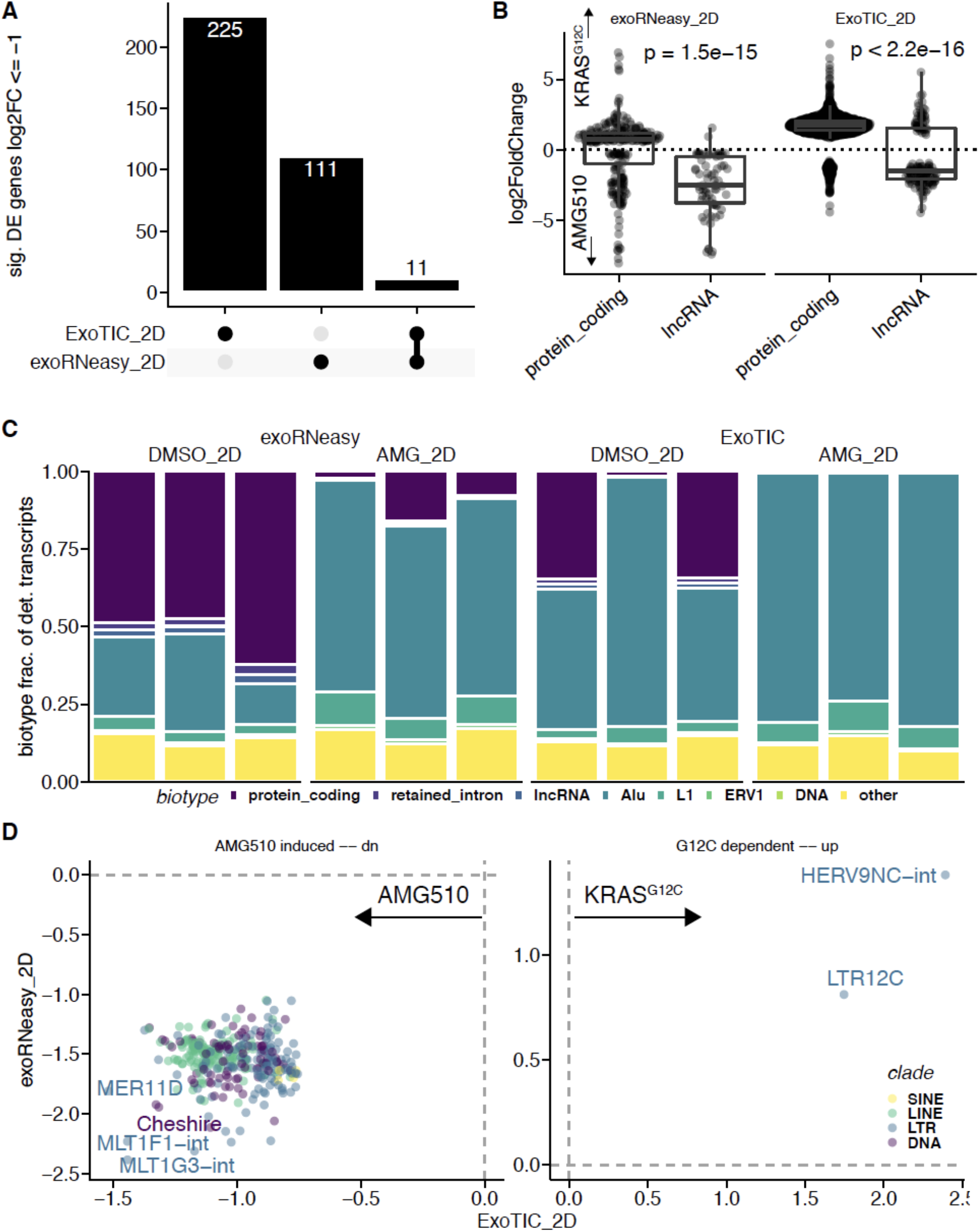
KRAS(G12C) inhibition increases secretion of noncoding RNAs in EVs. **A)** Upset plot of significantly downregulated genes (<=-1 log2FoldChange) in exoRNeasy- and ExoTIC-isolated EVs. **B**) Distribution of log2-scale fold-changes of genes from GENCODE protein-coding and lncRNA biotypes. **C**) Fraction of detected transcripts belonging to displayed biotypes and TE families. **D**) Scatter plot comparing log2-scale fold-changes of secreted TE RNAs isolated from control and AMG 510-treated H358 LUAD cells using the ExoTIC or exoRNeasy platforms.

To gain a more comprehensive understanding of how oncogenic KRAS(G12C) signaling affects the noncoding transcriptome, we next examined the TE RNA composition in EVs. TE RNAs are among the most abundantly expressed transcripts in the human genome^26^, and TE dysregulation has been observed in numerous cancers^27,28^. Moreover, oncogenic KRAS signaling specifically leads to TE RNA dysregulation through epigenomic reprogramming^8^. To determine the dynamics of TE RNA secretion in the context of KRAS(G12C) signaling, we included TE annotations in our RNA-seq analysis. The addition of TE annotations revealed that TE RNAs were the most abundant biotype in EVs, with the vast majority of exRNAs secreted from KRAS(G12C) inhibitor-treated cells being TE RNAs (**Fig. 3C**). Both EV isolation platforms revealed a common set of TE RNAs that was preferentially secreted in cells treated with KRAS(G12C) inhibitor (**Fig. 3D**).

### exRNA signatures of mutant KRAS(G12C) LUAD with prognostic potential

To determine the clinical relevance of common exRNA signatures identified using both the ExoTIC and exoRNeasy platforms, we examined RNA-seq data from the TCGA LUAD cohort. GSEA and DE analysis revealed the significant enrichment of 3 hallmark gene sets (MYC TARGETS V1, E2F TARGETS, and G2M CHECKPOINT) (**Fig. 4A**) and also a consensus set of 20 genes in mutant KRAS(G12C) LUAD samples across both our in vitro exRNA and in vivo TCGA datasets (**Fig. 4B**). Hierarchical clustering of TCGA LUAD patient RNA-seq data using the consensus 20-gene signature we identified from our exRNA data produced robust separation between mutant KRAS(G12C) LUAD tumor samples and healthy (WT) lung tissue samples (**Fig. 4C**). Furthermore, the average persample expression of this consensus signature was used to represent a patient ‘score’ for mutant KRAS(G12C) that produced significantly different distributions in KRAS(G12C) and WT samples, respectively (**Fig. 4D**). Lastly, the per-sample average of the KRAS(G12C) score was used to stratify the LUAD cohort into thirds, from which the top-third and bottom-third samples were used for Kaplan-Meier survival analysis. Patients with the highest KRAS(G12C) score (top-third) exhibited a significant decrease in survival probability (**Fig. 4E**), suggesting that our consensus 20-gene exRNA signature of mutant KRAS(G12C) signaling could potentially be used to predict clinical outcomes.

**Fig. 4.**
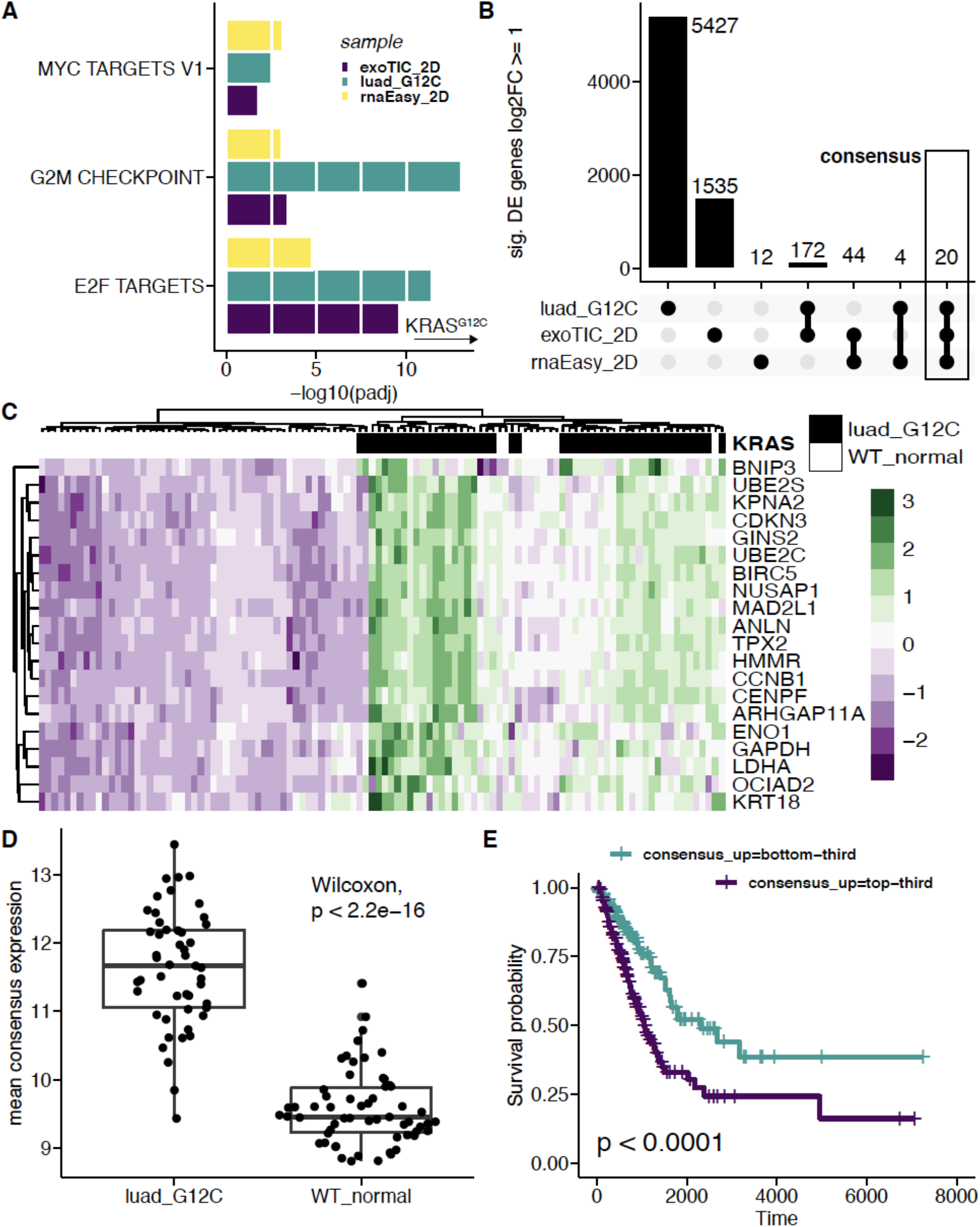
exRNA signatures of mutant KRAS(G12C) LUAD with prognostic potential. **A**) Bar plot of -log10 transformed adjusted p-values produced for each Hallmark gene set in Gene Set Enrichment Analysis across exoRNeasy, ExoTIC, and TCGA LUAD data sets. **B**) Upset plot quantifying overlap of upregulated genes (log2 fold-change >= 1) in exoRNeasy, ExoTIC, and TCGA LUAD differential expression. The labelled consensus set is used in the following panels. **C**) Heatmap with hierarchical clustering of scaled and centered count values for the 20 genes contained in the consensus overlapping set observed in **B**. **D**) Distribution of average expression of the consensus overlapping gene set in TCGA LUAD samples. Comparison of means with Wilcox demonstrates significant difference between the KRAS(G12C) tumor samples and the WT matched normal samples. **E**) Kaplan-Meier survival curve using overall survival of TCGA LUAD patients in the top-third and bottom third of consensus overlapping gene set expression.

## DISCUSSION

In this work, we identify mutant KRAS(G12C)-specific exRNA signatures from LUAD cells by comprehensively analyzing the coding and noncoding RNAs secreted in EVs. Our studies also demonstrate the value of using complementary affinity- and nanofiltration-based EV isolation methods for characterizing exRNAs. exRNAs reflect biological processes regulated by mutant KRAS(G12C), suggesting that they faithfully recapitulate aspects of intracellular gene expression dynamics in response to alterations in oncogenic KRAS signaling. Notably, exRNAs enriched upon KRAS(G12C) inhibition are significantly more variable, with strong and abundant noncoding RNA signals from both lncRNAs and TE RNAs. Moreover, KRAS(G12C)-dependent exRNA signatures detected across both EV isolation platforms reflect the in vivo RNA signatures in mutant KRAS(G12C) LUAD tumors from TCGA.

We also find that exRNA landscapes vary with differences in EV size. We sought to investigate two orthogonal approaches for EV isolation, which yielded distinct EV populations with variable size and exRNA content. These results highlight the importance of upstream EV isolation procedures prior to exRNA sequencing. Nanofiltration-based EV isolation was performed using the ExoTIC platform, which is a modular, size-based EV isolation tool that enables the capture of EVs centered around 155 nm and 120 nm from cells with and without intact KRAS(G12C) signaling, respectively^21^. The size distribution of EVs captured by ExoTIC overlaps significantly with the expected sizes of exosomes (30-150 nm), a subtype of small EVs secreted by most cell types and implicated as carriers of potential cancer biomarkers^10^. Furthermore, EV sizes decrease upon KRAS(G12C) inhibition, suggesting a potential link between KRAS and the regulation of EV size.

Consistent with our previous work^8^, noncoding RNAs are strongly enriched in EVs. We also observed a pronounced increase in library complexity upon the inclusion of TE annotations, with a significant amount of TE RNAs present in the secreted exRNA population. Although canonically silenced, TEs become activated in certain cancers and contribute to pathological events in these malignancies^27,29–32^. Our findings suggest that modulating KRAS signaling may also affect a subset of TEs. Two TEs in particular were enriched in both EV contexts: HERV9NC-int and its associated long terminal repeat (LTR), LTR12C^33^. This suggests a KRAS-dependent activation of this LTR promoter, which agrees with previous observations of mutant KRAS- and mutant TP53-mediated LTR activation^34–36^.

Finally, we explored the potential utility of KRAS(G12C)-specific exRNAs as predictive biomarkers for LUAD patient clinical outcomes. We evaluated our exRNA findings with the TCGA databases for KRAS(G12C) LUAD and healthy lung. Consensus genes that were enriched across exoRNeasy EVs, ExoTIC EVs, and TCGA LUAD datasets revealed a 20-gene signature capable of clustering KRAS(G12C) LUAD samples from their healthy counterparts. We find that elevated expression of this consensus 20-gene panel is associated with significant reduction in overall survival probability in LUAD patients. Our results reveal mutant KRAS(G12C)-specific exRNA signatures that may serve as diagnostic and prognostic biomarkers of lung cancer.

## MATERIALS AND METHODS

### Cell Lines

H358 lung adenocarcinoma cell lines harboring a KRAS(G12C) mutation were cultured in RPMI 1640 medium (Invitrogen) supplemented with 10% fetal bovine serum (Sigma) at 37°C, 5% CO2 in a humidified incubator. All cell lines tested negative for mycoplasma. The cell lines were purchased from American Type Culture Collection (ATCC).

### Cell viability assays

For adherent viability assays, 2.5E+04 cells/well were seeded in 96-well plates and incubated at 37°C, 5% CO2 for 16 hours. Then serially-diluted AMG 510 and DMSO were added to the cells, and plates were incubated in standard culture conditions for 72 hours. Cell viability was measured using a CellTiter-Glo® Luminescent Cell Viability Assay kit (Promega) according to the manufacturer’s protocol. The luminescence signal of treated samples was normalized to DMSO control. For spheroid viability assays, 5.0E+04 cells/wells were seeded in individual ultra-low adhesion 96-well plates (Corning) and incubated at 37°C, 5% CO2 for 24 hours. The cells were grown in standard culture conditions for 4 days. They were then harvested, and ATP production was measured using the CellTiter-Glo® Luminescent Cell Viability Assay kit (Promega) following the manufacturer’s protocol. The luminescence was measured on a SpectraMax iD3 molecular devices.

### RNA Isolation & Purification

RNA was isolated from H358 cells using Quick-RNA Mini-Prep kit (Zymogen). All RNA was quantified via NanoDrop-8000 Spectrophotometer. exRNA was isolated from cell culture media by two methods: exoRNeasy (Qiagen) and Exosome Total Isolation Chip (ExoTIC). In both methods, conditioned media was centrifuged at 300xg 4C for 5 min to remove possible cell debris and 2000xg 4C for 20 min to remove contaminating larger vesicles. Extracted EV sizes were examined by nanoparticle tracking analysis (NanoSight LM10). Extracted exRNAs were quantified via Qubit RNA HS assay kit (Thermo) and the QuantiFluor RNA System (Promega). RNA quality was examined by an Agilent 2100 Bioanalyzer (Agilent).

### Extracellular RNA sequencing

RNA sequencing libraries were prepared from exRNAs isolated using the exoRNeasy and ExoTIC EV isolation methods from adherent monolayer H358 cells using SMART-Seq (Takara Bio) according to the manufacturer’s protocol. The AMPure XP PCR purification kit was used for clean up and size selection of cDNA and final libraries. cDNA and library quality were determined using the High Sensitivity DNA Kit on a Bioanalyzer 2100 (Agilent Technologies). Multiplexed RNA-seq libraries were sequenced as paired end 150 runs on a NextSeq 500 to a sequencing depth of ~5 million read pairs.

### RNA-seq analysis

exRNA reads were first trimmed with Trimmomatic (0.39) and quantified with Salmon (1.30) with an index created using version 35 of the GENCODE reference transcriptome. The resulting transcript counts were aggregated to the gene level with Tximport and normalized with DESeqII. TE annotations were based on the Hg38 repeat track hosted on the UCSC Genome Browser. TCGA LUAD counts and metadata were downloaded from the UCSC Xena Browser in a DESeqII-normalized format^37^.

### Differential Expression and Gene Set Enrichment Analysis

DESeqII was used to estimated differential expression in all contexts with a standard model employing a formula dependent only on condition (AMG vs. DMSO, tumor vs. normal): ~ condition. Input counts were filtered to contain genes with at least 10 total counts as determined by Salmon. DESeq output was filtered to results that had an adjusted p-value of at most 0.05. Shrunken log2 fold change values were sorted, scaled, and used to rank the differentially expressed genes as input to Gene Set Enrichment Analysis (GSEA) performed using the R package *fgsea* with the ‘eps’ argument set to 0.0. Gene sets were acquired from MsigDB using the R package *msigdbr.* GSEA results were filtered to an adjusted p-value of at most 0.05.

### Sample clustering and dimensionality reduction

Hierarchical clustering was performed with the R package *pheatmap* using scaled, centered DESeqII normalized counts. Principal component analysis was performed with the R pakage *prcomp* and utilized DESeqII normalized counts filtered to genes/TEs with a coefficient of variance greater than or equal to the median across the reference.

### Kaplan-Meier survival analysis

Kaplan-Meier analysis was performed with the R package *survival.* TCGA LUAD samples were stratified into thirds based on their average expression of the gene set of interest. These strata were compared for differences in Overall Survival using the *survfit* function from *survival* with default parameters.

### Statistical analysis

All statistical analyses were performed in the R programming language (4.0.2) provided in a Docker container by the Rocker Project. Wilcoxon tests were performed using the R function *stat_compare_means* which calls built-in R functions *wilcox.test* and *t.test,* respectively.

## ACKNOWLEDGEMENTS

We thank members of the Kim Lab and Demirci Lab for helpful discussions. This work was supported by funds from the Baskin School of Engineering (to D.H.K.), the Ken and Gloria Levy Fund for RNA Biology (to D.H.K.), and the Department of Defense Congressionally Directed Medical Research Program (W81XWH-20-1-0746) (to U.D. and D.H.K.). R.E.R. is supported by the National Institutes of Health (1F99DK131504-01) and D.C. is supported by the Tobacco-Related Disease Research Program (T30DT0997). Illustrations were created with BioRender.com.

## AUTHOR CONTRIBUTIONS

D.H.K. conceptualized research, R.K. and D.H.K. designed research, R.K., M.O., S.V.M., and D.C. performed research, R.K. and R.E.R. analyzed data, U.D. and D.H.K. supervised research, and R.K., R.E.R., and D.H.K. wrote the paper with input from all the authors.

## SUPPLEMENTARY FIGURES

**Supplementary Fig. 1.**
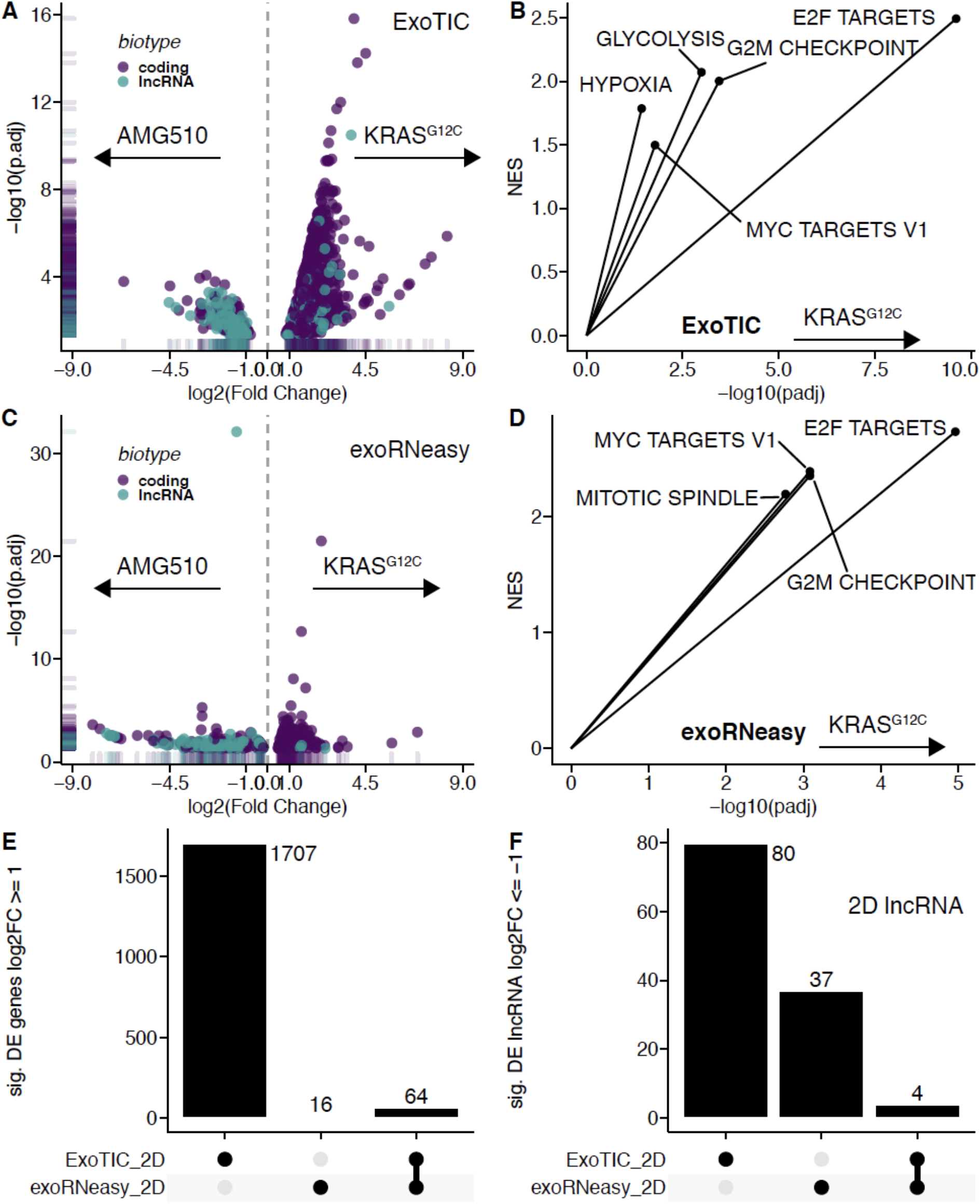
**A)** Overall differential expression volcano plot for ExoTIC-isolated exRNAs with and without AMG 510 treatment. **B)** Significantly enriched hallmark gene sets in ExoTIC-isolated exRNAs. **C)** Overall differential expression volcano plot for exoRNeasy-isolated exRNAs with and without AMG 510 treatment. **D)** Significantly enriched hallmark gene sets in exoRNeasy-isolated exRNAs. **E)** Upset plot of unique and overlapping significantly upregulated genes (>=1 log2FoldChange) in exoRNeasy- and ExoTIC-isolated exRNAs. **F)** Upset plot of unique and overlapping significantly downregulated lncRNA genes (<=-1 log2FoldChange) in exoRNeasy- and ExoTIC-isolated exRNAs.

